# Yellow fever virus is susceptible to sofosbuvir both *in vitro* and *in vivo*

**DOI:** 10.1101/266361

**Authors:** Caroline S. de Freitas, Luiza M. Higa, Carolina Sacramento, André C. Ferreira, Patrícia A. Reis, Rodrigo Delvecchio, Fabio Lima Monteiro, Giselle Barbosa-Lima, Yasmine Rangel Vieira, Mayara Mattos, Lucas Villas Bôas Hoelz, Rennan Papaleo Paes Leme, Mônica M. Bastos, Fernando A. Bozza, Patrícia T. Bozza, Nubia Boechat, Amilcar Tanuri, Thiago Moreno L. Souza

**Affiliations:** Laboratório de Imunofarmacologia, Instituto Oswaldo Cruz (IOC), Fundação Oswaldo Cruz (Fiocruz), Rio de Janeiro, RJ, Brazil.; Instituto de Biologia, Universidade Federal do Rio de Janeiro (UFRJ), Rio de Janeiro, RJ, Brazil.; Instituto Nacional de Infectologia (INI), Fiocruz, Rio de Janeiro, RJ, Brazil.; Instituto de Tecnologia de Fármacos (Farmanguinhos), Fiocruz, Rio de Janeiro, RJ, Brazil.; National Institute for Science and Technology on Innovation on Neglected Diseases (INCT/IDN), Center for Technological Development in Health (CDTS), Fiocruz, Rio de Janeiro, RJ, Brazil.

**Keywords:** Yellow fever virus, Yellow fever, antiviral, sofosbuvir

## Abstract

Yellow fever virus (YFV) is a member of the Flaviviridae family, that causes major mortality. In Brazil, YFV activity increased in the last years. It has been registered that sylvatic, instead of urban, yellow fever (YF) leads our contemporary public health concern. Low vaccinal coverage leaves the human population near the jangle vulnerable to the outbreak, making it necessary to identify therapeutic options. Repurposing of clinically approved antiviral drugs represents an alternative for such identification. Other Flaviviruses, such Zika (ZIKV) and dengue (DENV) viruses, are susceptible to Sofosbuvir, a clinically approved drug against hepatitis C virus (HCV). Moreover, sofosbuvir has a safety record on critically ill hepatic patients, making it an attractive option. Our data show that YFV RNA polymerase uses conserved amino acid resides for nucleotide binding to dock sofosbuvir. This drug inhibited YFV replication in different lineages of human hepatoma cells, Huh-7 and HepG2, with EC_50_ value of 4.8 µM. Sofosbuvir protected YFV-infected neonatal Swiss mice from mortality and weight loss. Our pre-clinical results indicate that sofosbuvir could represent an option against YFV.

## 1 Introduction

Yellow fever virus (YFV) is a single-strand positive-sense RNA virus which belongs to the Flaviviridae family. Yellow fever (YF) outbreaks were very common throughout the tropical world until the beginning of the 20^th^ century, when vaccination and vector control limited the urban virus circulation (1). Classically, sylvatic and urban cycles of YFV transmission occurs. Non-human primates are sylvatic reservoirs of jungle YFV and non-immunized humans entering the rain forest and those living at the ecotone (between preserved rain forest and urban area) are highly susceptible to severe YF, which may be transmitted by mosquitoes from *Haemagogus* and *Sabeties* genuses (2). The virus is usually brought to the urban setting by a viremic human who was infected in the jungle (2). The urban cycle involves transmission of the virus among humans by vectors like *Aedes* spp. mosquitoes (2).

In Brazil, an endemic country for YF, failed to vaccinate a large proportion of susceptible population. This scenario of low human vaccinal coverage along with increased sylvatic YFV activity in primates has been occurring in Brazil since 2016, leading to human cases of YF burst and between the second semester of 2016 and June 2017, 777 confirmed cases of YF with 261 attributed deaths and 1,659 epizootics reported (3). In fact, cases of YF increased 1.8-times compared to previous 35 years (3). Altogether these data show that unequivocally YF dramatic spread throughout Brazilian rain forests close to major cities in Southeastern region. Despite the detection of YFV in some urban areas, in humans and primates, during this recent reemergence, Brazilian Ministry of Health (MoH) argue this is a sylvatic cycle with no urban autochthonous transmission. Indeed, most of YFV recent activity was observed in areas adjacent to the Atlantic forest, where YFV genotype I was introduced two times, in 2005 (95% interval: 2002–2007) and 2016 (95% interval: 2012–2017), spilling over from the Amazon region (4).

YFV causes massive lethality for new world monkeys and around 30 % to humans (5). Once displaying severe signs of severe infection such as bleeding, shock, liver function decay and jaundice, infected individuals are likely to progress to poor clinical outcomes. Nevertheless, progression to acute hepatic failure is a reality for these individuals. No specific treatment option to YFV exist and patients solely receive just intensive palliative care.. Therefore, antivirals with anti-flavivirus activity may represent an important alternative for drug repurposing in an attempt to block or mitigate liver lesions induced by YFV replication.

Developed in the 1930s, YF 17D live-attenuated vaccine confers long-lasting immunity to its recipients. Vaccination is recommended for individuals aged ≥9 months who are living in or travelling to areas at risk for YF. Contraindications include hypersensitivity to vaccine components, severe immunodeficiency and age under 6 months–old (6). Even though 17D is highly effective and one of the safest vaccines in history, rare severe adverse events have been reported. YF vaccine-associated neurologic disease (YEL-AND) and YF vaccine-associated viscerotropic disease (YEL-AVD) which is similar to classic disease caused by wild-type YFV. Reporting rates of YEL-AND and YEL-AVD are 0.8 and 0.4 cases per 100,000 doses distributed (8). Specific treatment would be of utmost importance for individuals with yellow fever vaccine associated diseases and for YFV-exposed people for whom vaccination is contraindicated.

Our group has recently shown that sofosbuvir, a clinically approved anti-hepatitis C virus (HCV) drug, is also endowed with anti-Zika virus (ZIKV) antiviral activity in vitro and in vivo models (9, 10). Others have demonstrated that sofosbuvir also block dengue virus (DENV) replication (11). Sofosbuvir was approved by food and drug administration (FDA) in 2013. It has been used in therapy regimens thereafter in large worldwide scale to treat HCV-infected individuals, with infrequent registers of toxicity and adverse effects, even for complex patients, such as those co-infected with both HIV/HCV or with substantial liver damage (12-14). Sofosbuvir in very effective against against HCV genotype 1, 2 and 3, safety doses may range from 400 to 1200 mg daily for up to 24 weeks (12-14).

In the intention to have an active antiviral compound to inhibit YFV replication and block the progress of this disease to fatal outcomes, we tested whether sofosbuvir may possess anti-YFV activity. We found by *in silico* analysis that SFV binds to conserved amino acid residues on the YFV RNA RNA polymerase (NS5), inhibiting virus replication in human hepatoma cells and diminishing mortality in neonatal Swiss mice.

## 2 Material and Methods

### Reagents

The antivirals sofosbuvir and ribavirin were donated by the BMK Consortium and Instituto de Tecnologia de Farmacos (Farmanguinhos, Fiocruz), respectively. Drugs were dissolved in 100 % dimethylsulfoxide (DMSO) and subsequently diluted at least 10^4^-fold in culture or reaction medium before each assay. The final DMSO concentrations showed no cytotoxicity. The materials for cell culture were purchased from Thermo Scientific Life Sciences (Grand Island, NY) unless otherwise mentioned.

### Cells and Virus

Human hepatoma cell lines (Huh-7 and HepG2) and African green monkey (Vero) cells were cultured in DMEM supplemented with 10 % fetal bovine serum (FBS; HyClone, Logan, Utah), 100 U/mL penicillin, and 100 µg/mL streptomycin (15, 16). Cells were incubated at 37 °C in 5 % CO_2_.

YFV (vaccinal strain 17DD) was donated by the Reference Laboratory for Flavivirus, Fiocruz, Brazilian Ministry of Health. The virus was passaged at a multiplicity of infection (MOI) of 0.01 in Vero cells for 24 h at 37 °C. Virus titers were determined in Vero cell cultures by TCID_50_/mL (17).

### Cytotoxicity assay

Monolayers of 1.5 × 10^4^ Huh-7 cells in 96-well plates were treated for 5 days with various concentrations of sofosbuvir or ribavirin as a control. Then, 5 mg/ml 2,3-bis-(2-methoxy-4-nitro-5-sulfophenyl)-2*H*-tetrazolium-5-carboxanilide (XTT) in DMEM was added to the cells in the presence of 0.01 % of N-methyl dibenzopyrazine methyl sulfate (PMS). After incubating for 4 h at 37 °C, the plates were read in a spectrophotometer at 492 nm and 620 nm(18). The 50 % cytotoxic concentration (CC_50_) was calculated by a non-linear regression analysis of the dose-response curves.

### Yield-reduction assay

Monolayers of 5.5 × 10^6^ Huh-7 cells in 6-well plates were infected with YFV at MOI of 0.1 for 1 h at 37 °C. The cells were washed with PBS to remove residual viruses, and various concentrations of sofosbuvir, or ribavirin, in DMEM with 1 % FBS were added. After 24 h, the cells were lysed, the cellular debris was cleared by centrifugation, and the virus titers in the supernatant were determined as TCID_50_/mL in Vero cells. A non-linear regression analysis of the dose-response curves was performed to calculate the concentration at which each drug inhibited the plaque-forming activity of ZIKV by 50 % (EC_50_).

### Flow Cytometry

HUH7 and HepG2 cells were seeded in 6-well plates at density of 6 x10^4^/well and 2.5 × 10^5^ cells/well, respectively. For infection, the growth medium was replaced by serum-free medium containing the virus at MOI of 1. Mock-infected cells were incubated with conditioned media from uninfected cells prepared exactly as performed for viral propagation. After 1 hour, inoculum was removed and replaced by growth medium containing either the vehicle or 0.4, 2, 10 and 50 μM of sofosbuvir and incubated for 48 hours for HepG2 cells and 72 hours for HUH7 cells. After this period, cells were harvested by treatment with a 0.25% trypsin solution. Cells were fixed with 4% paraformaldehyde (Sigma Aldrich) in phosphate buffered saline (PBS) for 15 min at room temperature and washed with PBS. Cells were permeabilized with 0.1% Triton X-100 (Sigma Aldrich) in PBS, washed with PBS, and blocked with PBS with 5% FBS. Cells were incubated with 4G2, a pan-flavivirus antibody raised against the ZIKV envelope E protein produced in 4G2-4-15 hybridoma cells (ATCC), diluted 1:10 in PBS with 5% FBS. Cells were labeled with donkey anti-mouse Alexa Fluor 488 antibody (Thermo Scientific, Waltham, MA, USA) diluted 1:1000 in PBS with 5% FBS, and were analyzed by flow cytometry in a BD Accuri C6 (Becton, Dickinson and Company, Franklin Lakes, NJ, USA) for YFV infection.

### Sequence comparisons

The sequences encoding the C-terminal portion of the RNA polymerase from Flaviviruses were acquired from the complete sequences deposited in GenBank. An alignment was performed using the ClustalW algorithm in the Mega 6.0 software. The sequences were analyzed by disparity index per site. Compared regions are displayed in the supplementary File 1.

### Comparative modeling

The amino acid sequence encoding YFV RNA polymerase (YFRP; UniProtKB code: P03314) was obtained from the EXPASY proteomic portal(19) (http://ca.expasy.org/). The template search was performed using the Blast server (http://blast.ncbi.nlm.nih.gov/Blast.cgi) with the Protein Data Bank(20) (PDB; http://www.pdb.org/pdb/home/home.do) as the database and the default options. The T-COFFEE algorithm was used to generate the alignment between the amino acid sequences of the template proteins and YFRP. Subsequently, the construction of the YFRP complex was performed using MODELLER 9.19 software(21), which employs spatial restriction techniques based on the 3D-template structure. The preliminary model was refined in the same software, using three cycles of the default optimization protocol. The structural evaluation of the model was then performed using two independent algorithms in the SAVES server (http://nihserver.mbi.ucla.edu/SAVES_3/): PROCHECK software (22) (stereochemical quality analysis) and VERIFY 3D (23) (compatibility analysis between the 3D model and its own amino acid sequence by assigning a structural class based on its location and environment and by comparing the results with those of crystal structures).

### Animals

Swiss albino mice (*Mus musculus*) (pathogen-free) from the Oswaldo Cruz Foundation breeding unit (Instituto de Ciência e Tecnologia em Biomodelos; ICTB/Fiocruz) were used for these studies. The animals were kept at a constant temperature (25°C) with free access to chow and water in a 12-h light/dark cycle. The experimental laboratory received pregnant mice (at approximately the 14th gestational day) from the breeding unit. Pregnant mice were observed daily until delivery to accurately determine the postnatal day. We established a litter size of 10 animals for all experimental replicates.

The Animal Welfare Committee of the Oswaldo Cruz Foundation (CEUA/FIOCRUZ) approved and covered (license number L-016/2016) the experiments in this study. The procedures described in this study were in accordance with the local guidelines and guidelines published in the National Institutes of Health Guide for the Care and Use of Laboratory Animals. The study is reported in accordance with the ARRIVE guidelines for reporting experiments involving animals(24).

### Experimental infection and treatment

Three-day-old Swiss mice were infected intraperitoneally with 2 × 10^6^TCID_50_/mL of virus, unless otherwise mentioned. Treatments with sofosbuvir were carried out with sofosbuvir at 20 mg/kg/day administered intraperitoneally. Treatment started one day prior to infection (pretreatment). Animals were monitored daily for survival and weight gain.

If necessary to alleviate animal suffering, euthanasia was performed. The criteria were the following: i) differences in weight gain between infected and control groups >50%, ii) ataxia, iii) loss of gait reflex, iv) absence of righting reflex within 60 seconds, and v) separation, with no feeding, of moribund offspring by the female adult mouse.

### Statistical analysis

All assays were performed and codified by one professional. Subsequently, a different professional analyzed the results before the identification of the experimental groups. This approach was used to keep the pharmacological assays blind. All experiments were carried out at least three independent times, including technical replicates in each assay. The dose-response curves used to calculate the EC_50_ and CC_50_ values were generated by Prism GraphPad software 7.0. The equations to fit the best curve were generated based on R^2^ values ≥ 0.9. Fisher’s exact and ANOVA tests were also used, with *P* values <0.05 considered statistically significant. The significance of survival curves was evaluated using the Log-rank (Mantel-Cox) test. *P* values of 0.05 or less were considered statistically significant.

## 3 Results

### Prediction of the complex between Sofosbuvir triphosphate and YFRP using comparative modeling

We initially compared the disparities among region encoding the RDRP region of the contemporary Flaviviruses, DENV, ZIKV and YFV. The YFV RDRP shares the conserved domain for catalytic activity with the orthologues enzymes (Supplementary File 1 and Table 1).

**Table 1.** Estimate disparities index per site between amino acid sequences encoding the RDRP domain peptide located at last 680 amino acid of C-Terminal from contemporary Flaviviruses

| Order | GenBank # | Agent | Distance comparisons |  |  |  |  |
| --- | --- | --- | --- | --- | --- | --- | --- |
|  |  |  | 1 | 2 | 3 | 4 | 5 |
| 1 | FJ654700.1 | YFV |  |  |  |  |  |
| 2 | KX197205.1 | ZIKV | 0.000 |  |  |  |  |
| 3 | AB189124.1 | DENV1 | 0.000 | 0.002 |  |  |  |
| 4 | DQ672564.1 | DENV2 | 0.000 | 0.000 | 0.056 |  |  |
| 5 | AY858046.2 | DENV3 | 0.000 | 0.107 | 0.000 | 0.062 |  |
| 6 | GU289913.1 | DENV4 | 0.000 | 0.042 | 0.000 | 0.011 | 0.000 |

Next, the crystal structures of DENV NS5 complexed with viral RNA (PDB code: 5DTO) (25), HCV NS5B in complex with Sofosbuvir diphosphate (PDB code: 4WTG)(26), and complexed to UTP (PDB code: 1GX6) (27) were selected and used in the comparative modeling procedure, covering 100% of the YFRP sequence considered here (residues Thr252-Ile878). These three template proteins represent orthologous viral RNA RNA polymerases from the Flaviviridae family. Consequently, the resulting 3D model of YFRP in complex with Sofosbuvir triphosphate showed good structural quality.

The analysis of YFRP model suggests that Sofosbuvir triphosphate binds between the palm and the fingers regions of YFRP, making hydrogen bonds with Gly538, Trp539, Ser603, and Lys 693 residues and salt bridge interactions with Lys359 and two Mg^2+^ ions. Interestingly, these interactions are all described as relevant for incorporation of natural ribonucleotides (26) (Figure 1). Therefore, these results motivate further testing in biologically relevant models to YFV replication.

**Figure 1.**
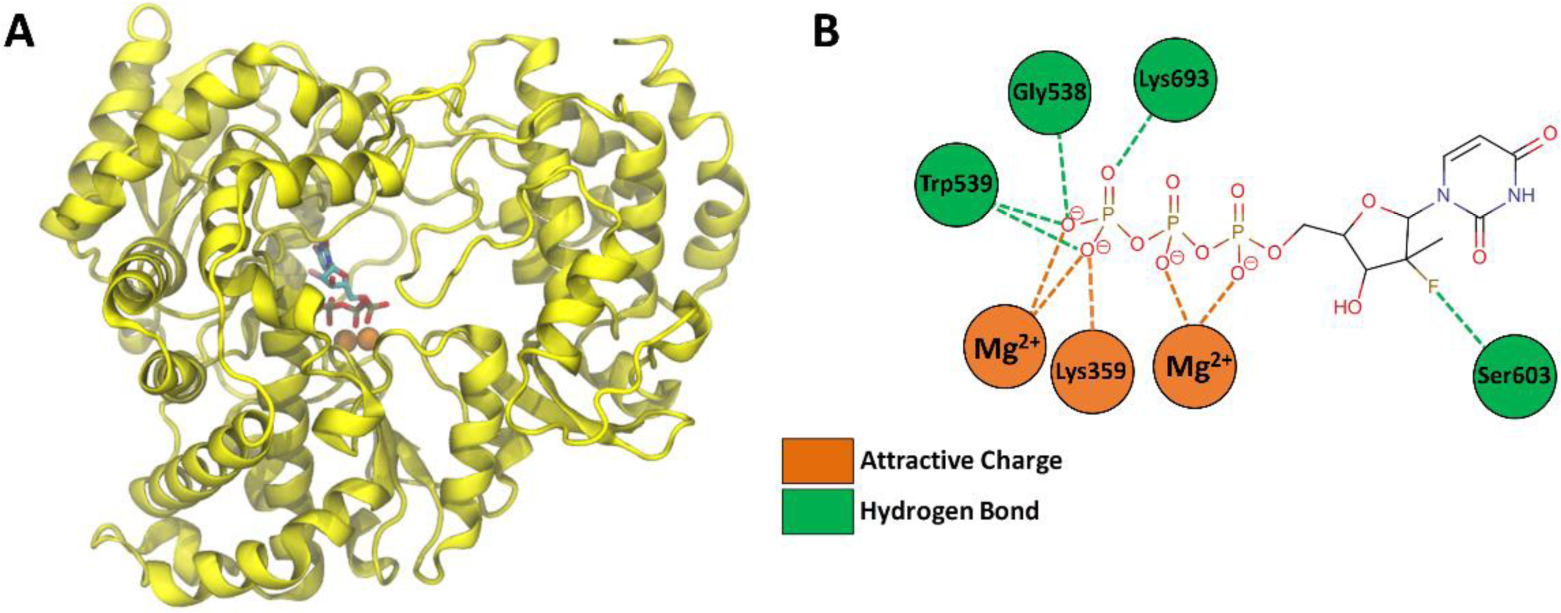
3D model of YFV RNA polymerase (YFRP) in complex with Sofosbuvir triphosphate. Cartoon representation of YFRP 3D model. The Sofobuvir triphosphate structure is represented by sticks and colored according to atom type (red, oxygen; blue, nitrogen; orange, carbon; yellow, phosphorus) and the two catalytic Mg^2+^ ions are shown as orange spheres. (B) Schematic representation of the interaction between amino acid residues of YFRP and the sofosbuvir triphosphate structure, where the salt bridges are shown in orange and hydrogen bonds are in green.

### Sofosbuvir inhibits YFV replication in different models of human hepatocytes

YFV primarily replicates in the liver, where sofosbuvir is majorly converted from prodrug to its active metabolite (12-14). Thus, we monitored YFV susceptibility to sofosbuvir using human hepatocellular carcinoma cells. YFV was yield in Huh-7 cells in the presence of sofosbuvir for 24 h, when supernatant was harvested and tittered in Vero cells. Indeed, sofosbuvir inhibited in dose-dependent manner YFV replication (Figure 2A). As a control, ribavirin, a broad spectrum antiviral, also inhibited YFV replication (Figure 2A). Sofosbuvir’s and ribavirin’s potencies to inhibit YFV replication were quite comparable, with EC_50_ values equal to 4.8 ± 0.2 and 3.9 ± 0.3 µM (Table 1), respectively. Since sofosbuvir is around 25 % less cytotoxic than ribavirin – its selectivity index (SI), the ratio between EC_50_ and CC_50_, was higher (Table 2).

**Figure 2.**
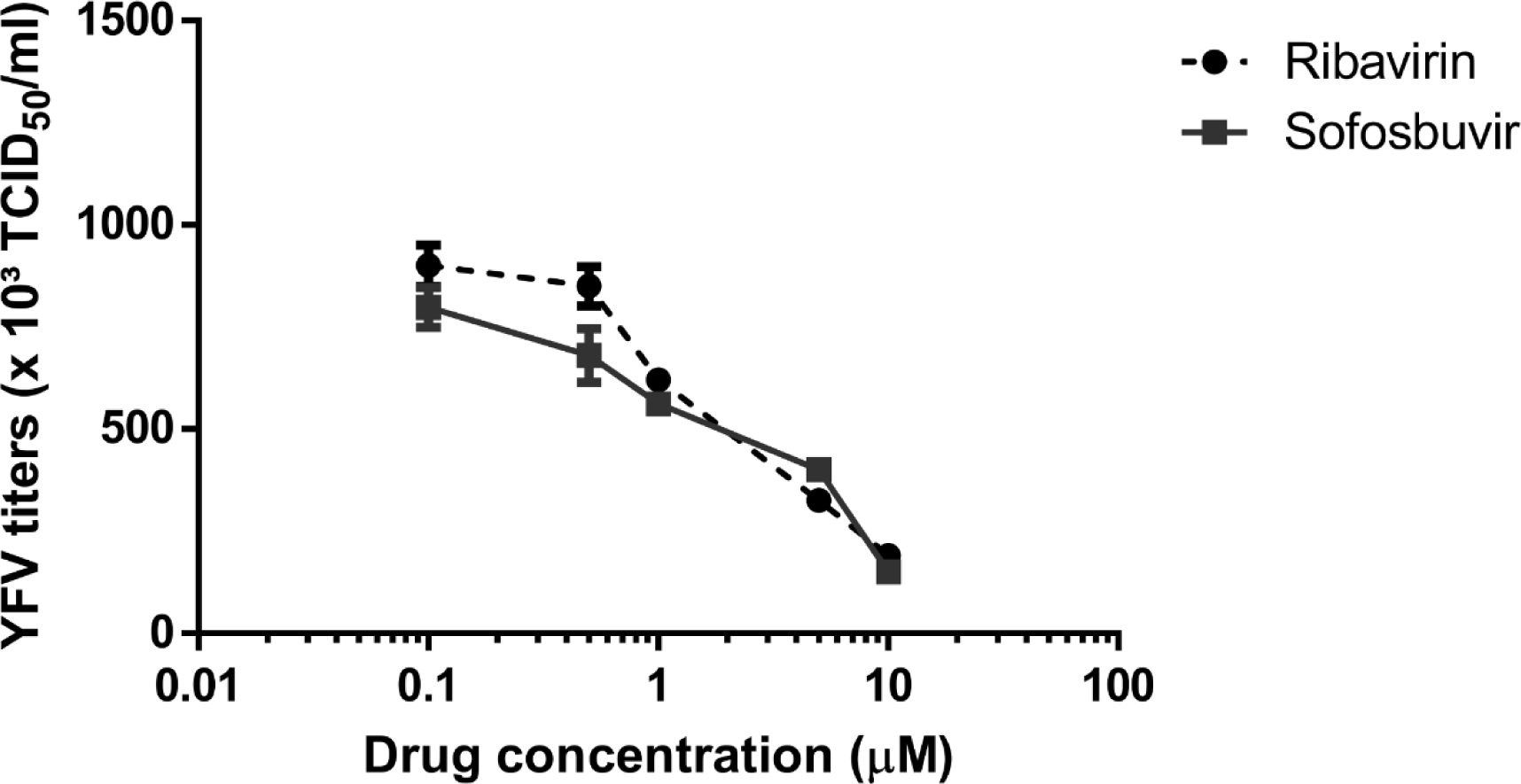
Pharmacology of sofosbuvir against YFV. Huh-7 were infected with YFV at MOI of 0.1 and exposed to various concentrations of sofosbuvir or ribavirin (B) for 24 h. Supernatant was harvested and tittered in Vero cells by TCID_50_/mL. The data represent means ± SEM of three independent experiments.

**Table 2.** Pharmacological parameters related to antiviral activity of sofosbuvir and ribavirin against YFV

| Drugs | EC<sub>50</sub> | CC<sub>50</sub> | SI* |
| --- | --- | --- | --- |
| Sofosbuvir | 4.8 ± 0.2 | 381 ± 25 | 80 |
| Ribavirin | 3.9 ± 0.3 | 284 ± 12 | 72 |

\*Selectivity index (SI) = CC<sub>50</sub>/EC<sub>50</sub>

In order to gain further knowledge about the antiviral properties of sofosbuvir in hepatic cells, viral infectivity was assessed by flow cytometry using the 4G2 antibody, a pan-flavivirus antibody raised against the envelope protein (Fig 3). Huh7 cells were infected with YFV at a MOI of 1 and then treated with sofosbuvir in concentrations ranging from 0.4 to 50 μM for 72 hours. Sofosbuvir dramatically reduced the number of YFV-infected cells in a dose-dependent manner. Doses above 2 µM totally abolished YFV antigen production in Huh7 infected cells (Fig 3A). Additionally, sofosbuvir treatment resulted in decreased production of infectious virus particles (Fig 3A) also in a dose-dependent manner. Treatment with 10 μM and 50 μM sofosbuvir led to a 4,034 and 561,142-fold reduction in the level of infectious particles. Similar results were obtained for HepG2 cells (Fig 3B). Sofosbuvir treatment resulted in a reduction in the number of YFV-infected cells and in the detection of infectious YFV (Fig 3B) compared to untreated cells. These results indicate that sofosbuvir is a good candidate against YF, deserving further *in vivo* testing.

**Figure 3.**
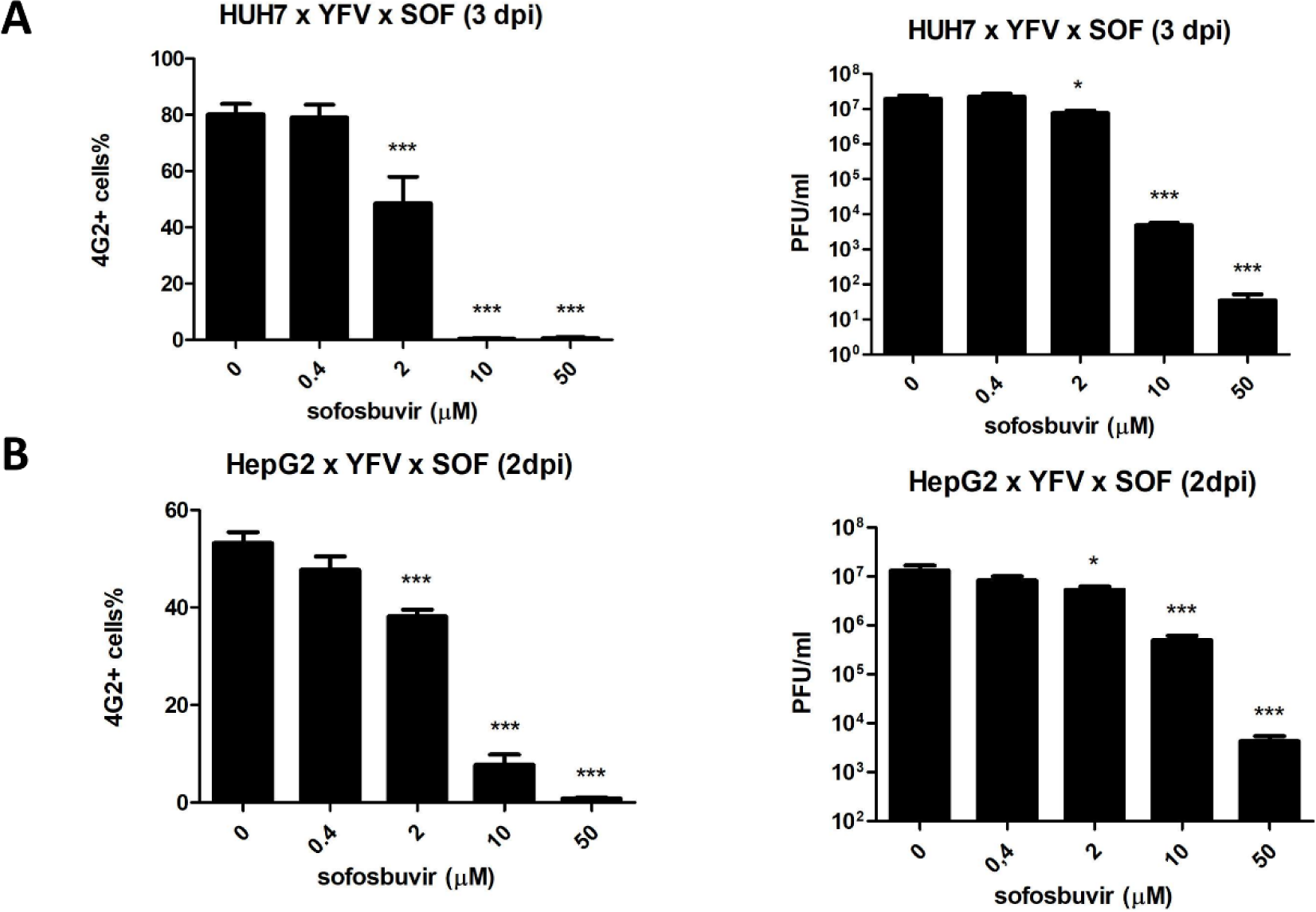
Sofosbuvir inhibits YFV protein production. Huh-7 (A) or HepG2 (B) were infected at MOI of 0.1 and exposed to key concentration of sofosbuvir for 3 or 2 days, respectively. The time points in this experiment were adjust to observe maximum protein labeling. Anti-pan-flavivirus antibody was used for cytometric analysis (left panels); in parallel, plaque forming unites were monitored (right panels). The data represent means ± SEM of three independent experiments. * P < 0.05

### Sofosbuvir enhances survival of YFV-infected neonatal mice

To test sofosbuvir *in vivo*, we used the dose of 20 mg/kg/day in newborn Swiss outbred mice. This dose is consistent with its pre-clinical/clinical studies for drug approval(14). Sofosbuvir protected treated mice from YFV-induced mortality (Fig 4A). Infected mice died within two weeks after infection, whereas 70% of sofosbuvir-treated YFV-infected animals survived to lethal inoculum (Fig. 4A). We also evaluated the weight gain during the time course of our experiment in mice. YFV-infected animals had reduced postnatal development, whereas sofosbuvir-treated YFV-infected mice gained weight almost as much as the uninfected controls (Fig. 4B).

**Figure 4.**
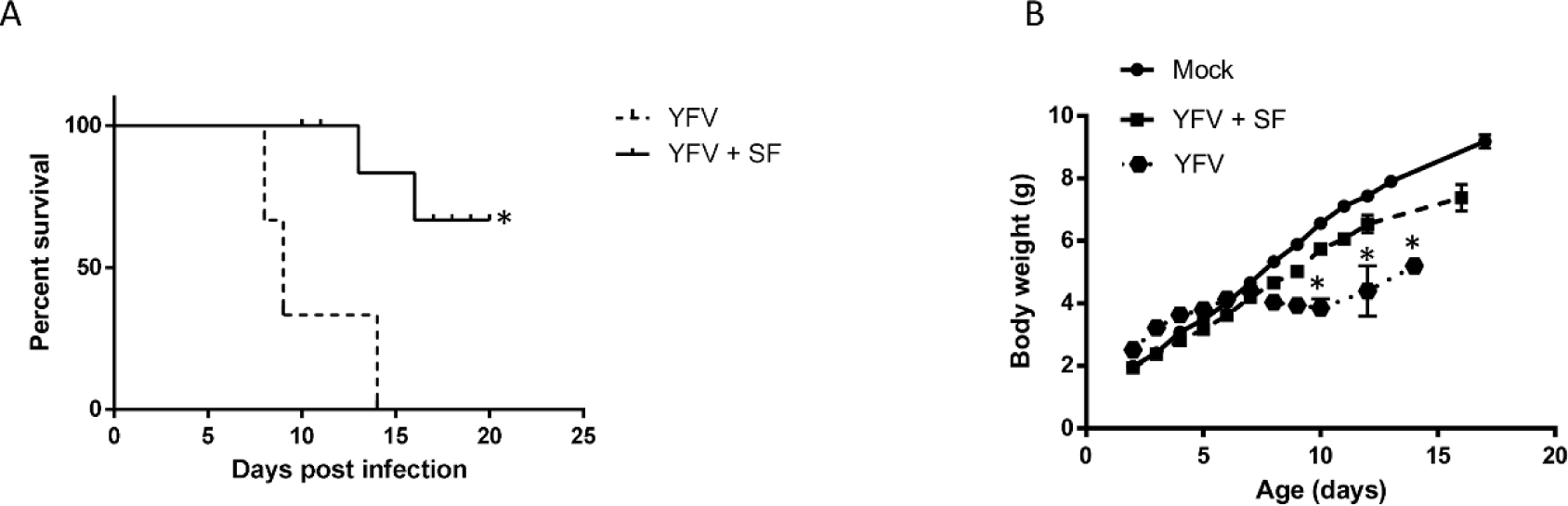
Treatment with Sofosbuvir increases survival and inhibits weight loss of YFV-infected mice. Three**-**day-old Swiss mice were infected with YFV (2 × 10^6^ TCID_50_) and treated with sofosbuvir either 1 day before. Survival (A) and weight variation (B) were assessed during the course of treatment. Survival was statistically assessed by Log-rank (Mentel-Cox) test. Differences in weight are displayed as the means ± SEM, and twoway ANOVA for each day was used to assess the significance. Independent experiments were performed with at least 10 mice/group. * P < 0.05.

## 4 Discussion

Brazil has been challenged in recent years by the (re)emergence of arboviruses. Massive cases of microcephaly associated with ZIKV circulation were registered in 2015/16 (28). Two different genotypes of chikungunya virus (CHIKV) co-circulate since 2014 (29, 30). The four DENV subtypes are hyperendemic throughout the country (31). More recently, YFV activity increased, affecting both non-human primates and humans without constituted immunity (3, 5). These facts clearly demonstrate that strategies of vector control failed and highlights the necessity of other ways to control or even mitigate the diseases provoked by the arboviruses.

In the context of this article, re-emergence of YFV also points out that poor vaccination coverage left human population living or entering the ecotone susceptible to this virus. Unvaccinated individuals may quickly progress to severe YF, with hepatic and even neurological impairment (1). In addition, due to massive YF vaccine captains a fair amount of vaccine related severe adverse events may be expected. Thus, finding antiviral drugs to treat YFV-infected individuals is critical for medical intervention in critical cases.

Several groups demonstrated that the clinically approved anti-HCV drug sofosbuvir is endowed with antiviral activity towards other flavivirus such as ZIKV (9, 10) and DENV (11). Considering that these agents belong to the same family of viruses, it was plausible to test sofosbuvir against YFV.Indeed, we observed that sofosbuvir predictively targets the YFV RNA RNA polymerase *in silico* and by dose response antiviral assays. In fact, the *in vitro* antiviral results display that sofosbuvir’s EC50% towards YFV is in micromolar range and comparable to what has been described for ZIKV and DENV (9-11). Sofosbuvir’s metabolism is complex and some facets are clearly undescribed. Although it was thought to be converted from prodrug to active compound only on liver cells, we found it also happens on human neural stem cells (10). Given this complexity, *in vivo* testing of sofosbuvir is pivotal to better comprehend its pharmacology and *in vivo* antiviral potency.

Our data showed that sofosbuvir inhibits YFV in hepatic cell lines. This is particularly relevant since YFV targets hepatocytes and the liver is the most affected organ in YF (32). The degree of liver damage measured by elevated aspartate aminotransferase and jaundice is associated with higher mortality (33). Massive apoptosis and necrosis of hepatocytes are reported in fatal cases (34). Additionally, impaired synthesis of clotting factors caused by YFV-induced liver injury is key to the pathogenesis of haemorrhagic manifestations in severe YF (35). Therefore, sofosbuvir treatment may improve liver function in YF patients and hopefully impact clinical outcome.

In line with our results, sofosbuvir reduced the YFV-induced mortality and lack of weight gain in neonatal mice. Considering our data and the safety history of sofosbuvir in hepatitis, it is not a hyperbolic conclusion to consider sofosbuvir in the clinical use in YF infection in humans. Primarily, sofosbuvir could be worthwhile for acutely infected individuals and those displaying

YF vaccine associated neurotropic and viscerotropic diseases provoked by virus replication. Since vaccine is not recommended by especial groups of individuals at higher risk of severe adverse events, such as elderly and those with immunedeficiencies, sofosbuvir could be used prophylactically. The results described here originally demonstrates the antiviral activity of sofosbuvir to YFV, which is causing an ongoing epidemic in Brazil, providing the primary scientific evidence for a new use of a clinically approved antiviral drug.

## Acknowledgments

Thanks are due to Dr. Karin Brüning, from the BMK Consortium (Blanver Farmoquímica Ltda; Microbiológica Química e FarmacêuticaLtda; Karin Bruning & Cia. Ltda), for donating sofosbuvir, and to Dr Ana M. B. de Filippis, from Laboratório de Flavivírus, Instituto Oswaldo Cruz, Fiocruz, for kindly provide the YFV strain.

## Author contributions

Experimental execution and analysis - CSF, LMH, CS, ACF, PAR, RD, FL, GBL, YRV, MM, LVBH, RPPL

Data analysis, manuscript preparation, and revision – CSF, LMH, CS, ACF, PAR, RD, FL, MMB, FAB, PTB, NB, AT, TMLS

Conceptualized the study – AT, TMLS

All authors revised and approved the manuscript.

## Additional information

**Role of funding:** This work was supported by Conselho Nacional de Desenvolvimento e Pesquisa (CNPq), Fundação de Amparo à Pesquisa do Estado do Rio de Janeiro (FAPERJ).

